# TAXISCAN, Optimizing throughput and behavioral depth in standard *C. elegans* chemotaxis assays

**DOI:** 10.64898/2026.07.21.739905

**Authors:** Victoria R. Yarmey, Daisy Aguilar Aguilar, Adriana San-Miguel

## Abstract

Chemosensory signaling is crucial for organisms to respond to their environment and dysfunctions in these complex pathways have been implicated in diseases such as neurological disorders. The nematode *C. elegans* is a useful tool for studying chemosensation as it offers a fully-mapped nervous system with relatively simple chemosensory circuits. One of the most common methods to quantify chemosensory function in worms is through chemotaxis assays, which require extensive time and manual effort. In this work, we sought to capture novel chemosensory metrics and increase the throughput of these assays by combining rapid image acquisition with bioinformatics processing. In doing so, we have developed a Technique for Automating chemotaXis Investigations using Simple-Capture Approaches in Nematodes (TAXISCAN), a rapid acquisition platform that can be used to study natural chemosensory behavior as well as to characterize behavioral and locomotion phenotypes in half the time of standard assays.

**Highlights:**

- The TAXISCAN platform provides a rapid, accurate, and inexpensive means of conducting unbiased chemotaxis assays in *C. elegans*
- In addition to the Chemotaxis Index (CI), the TAXISCAN platform can quantify behavioral metrics such as movement speed, distance travelled, and dispersion patterns
- We successfully use TAXISCAN to discriminate behavioral phenotypes in neurosensory mutants

## Introduction

The ability to detect and interpret chemical cues from the environment is a core survival strategy for finding food, avoiding danger, and communicating with other organisms. *Caenorhabditis elegans* (*C. elegans*) is a widely used model organism for studying chemosensory circuits as it possesses a fully mapped nervous system with a well-characterized panel of responses to various chemical stimuli [1–3]. These responses include chemotaxis, a crucial survival tactic involving the movement towards or away from both volatile (olfactory) and water-soluble (gustatory) stimuli [1]. While this behavior protects the worm from danger and allows it to seek food and mates, it also reflects sensory function at the molecular, cellular, and physiological levels. Chemotaxis assays are thus a core tool for studying chemosensory function [4–10], organismal health [11,12], behavior, neuroplasticity, and memory [12–14].

The first *C. elegans* chemotaxis assay is attributed to Ward et al. in 1973 [15], and involved tracking worm movement across an agar plate in response to several chemoattractants (cyclic nucleotides, salts, alkaline substances, and amino acids). Despite several variations, the standard chemotaxis assay involves placing worms at the center of an agar plate, with a chemical stimulus (test compound) added to one side of the plate and its buffer added to the opposite side (S1 Fig). Worms are typically allowed 60 minutes to migrate across the plate, at which point the observer counts the number of worms on either side. The Chemotaxis Index (CI) measures the fraction of worms that moved towards (+1) or away from (−1) the test compound, thus representing the general attraction or avoidance behaviors of the entire plate. Many studies divide chemotaxis plates into halves [2,11,16,17], though several have also opted for quadrants [18–20], or gradient fields [21–24]; the resulting CI is dependent on its respective plate configuration. Many chemotaxis assays incorporate an immobilization step to prevent worms from migrating between regions during scoring, which can be accomplished by temporarily placing the plates at 4 °C [1,3,18]. Since *C. elegans* is capable of rapid decision-making and learning, enabling it to respond differently to the same stimulus over time, it is also common to use a paralytic like sodium azide (NaN_3_) that gradually diffuses across the agar to capture the initial behavioral response to the stimulus [25,26].

Several factors impact chemotaxis assays, leading to high day-to-day variability and issues with reproducibility [2,18]. These include the health, age, and genetic background of the worm [3,6,11,27–29] but also minute changes in the temperature, humidity, sound, light, and the presence of external odors in the laboratory environment [2,18,30]. Worm behavior is also affected by plate configuration, incubation time, chemical stimuli, and sample population sizes. Several attempts have been made to account for these factors through experimental design [2,12,18,24], but these assays still require large sample sizes to overcome their high biological variability. Traditional chemotaxis assays are labor-intensive, time-consuming, and associated with limited experimental throughput due to the manual handling and manual scoring of individual assay plates (consisting of 100-200 worms). To overcome this obstacle, approaches for single-plate tracking have been developed [27,31–33], where the movement of individual worms is automatically tracked by combining time-lapse videos with automated image analysis. Automated worm tracking can also capture additional behavioral metrics such as position, speed, distance, body bends, reversals, and pirouetting, which are not visible through CI. Platforms for simultaneous analysis of multiple populations [28,34] and microfluidic approaches [35–37] improve the throughput and detail of the assays, but require additional equipment or the fabrication of custom behavioral arenas [35–37].

In 2013, Stroustrup et al. introduced the *C. elegans* Lifespan Machine [36], an accessible platform that uses a document scanner to monitor population survival by measuring movement at an individual level across multiple plates simultaneously. Over a decade later, Fryer et al. adapted this capture platform to conduct large-scale chemotactic screens [34]. However, their method relies on custom-fabricated behavioral arenas and a liquid handler, which, while greatly improving throughput and accuracy, limits the platform’s utility. In this work, we developed TAXISCAN (a Technique for Automating chemotaXis Investigations using Simple-Capture Approaches in Nematodes), a platform adapted from the Lifespan Machine for automated and in-depth chemotaxis analysis without requiring additional equipment. TAXISCAN simultaneously captures images of multiple standard chemotaxis plates and automatically determines individual worm position relative to the chemical stimuli. Using individual position information, TAXISCAN enhances chemotaxis analysis by expanding the characterization of behavioral and locomotion phenotypes in *C. elegans* beyond their primary chemosensory responses. We further adapted this platform to enable time-lapse recording and extract additional metrics descriptive of the population-level migration path. In addition to simplifying and accelerating the CI scoring, TAXISCAN significantly improved the throughput and depth of the standard chemotaxis assays enabling the analysis on large scales.

## Materials & Methods

### *C. elegans* strains & maintenance

Animals were maintained at 20 °C on nematode growth medium (NGM) agar plates supplemented with OP50 *E. coli.* All experiments were conducted with populations that had been well-fed for at least 3 generations. To age-synchronize worms for experiments, eggs were collected from gravid adults via bleaching [38] (0.5 M NaOH, 1.25% Sodium Hypochlorite) with 3 washes of M9 Buffer with 0.01% v/v Triton-X100 (M9+TX). All strains used in this study were acquired from the *Caenorhabditis* Genetics Center (CGC, University of Minnesota). (Table 1).

**Table 1.** Strains used in this study.

| Strain | Genotype |
| --- | --- |
| N2 | Wildtype Bristol |
| JT6924 | *daf-19(m86); daf-12(sa204)* |
| CX3933 | *kyls140 [str-2::GFP + lin-15(+)] ; slo-1(ky389)* |
| JPS428 | *slo-1(gk602291)* |
| NM1968 | *slo-1(js379)* |

### Chemotaxis assay preparation

To prepare assay plates, 5 cm flat-bottomed chemotaxis plates (2% agar, 1 mM CaCl_2_, 1 mM MgSO4, 5 mM K_2_PO_4_) were poured 5 days in advance and allowed to dry for 2 days at room temperature (RT) before storing at 4 °C. A stencil was used to mark the bottom of the plates in quadrant-plate configurations [3,18]: a thick dot at the center of the plate, a small dot approximately 1cm from the edge of quadrants QII and QIII, and a thick “X” approximately 1 cm from the edge of quadrants QI and QIV (S1 Fig). These markings indicate locations for the worms, 2 μL of the control compound, and 2 μL of the test compound, respectively. No other markings were used on the bottom of the plates to discourage errors with downstream image processing. CuSO_4_ was added to the plates 24 hr prior to running the assay while the volatile compounds Isoamyl alcohol (IAA) and 1-octanol, were added immediately before [19] or after [18] placing the worms. On the day of the assay, the plates were brought to RT for 10 minutes prior to adding 1 μL of 0.5 M NaN_3_ to the dots and “X’s” in the quadrants acting as paralyzing agent.

To conduct the chemotaxis assays, young adult (YA) worms were gently washed four times in chemotaxis buffer (1 mM CaCl_2_, 1 mM MgSO_4_, 5 mM K_2_PO_4_) with 0.01% v/v Triton-X100 (CB+TX) using gravity to pellet while restricting the entire wash process to under 20 minutes to avoid the effects of starvation [2]. Approximately 100-200 worms (2-5 μL) were added to the center of each chemotaxis plate, after which 2 μL of the volatile compounds and their buffers were added to their respective regions. Worms were exposed to chemotaxis buffer (Null) as a negative control, 1% IAA in 50% ethanol as an attractant, 10% 1-octanol in 50% ethanol as a volatile repellent, or 100 mM CuSO_4_ as a water-soluble repellent. The plates were covered and allowed to incubate in the dark at RT for 1 hr before moving to 4 °C for a minimum of 5 min to await scoring. To conduct standard scoring, an observer used a benchtop microscope to count the number of worms within every region of each plate. Image-scoring was conducted by scanning a plate and using the point tool in ImageJ to manually locate the approximate center of each worm in the image, then CI was determined by mapping the resulting coordinates to the assay quadrants. For live-scanning, the chemotaxis plates were prepared directly on the Epson V600 scanner bed, and images were taken at 5 min intervals for 60 min without transferring plates to 4 °C.

### Image acquisition

The bottoms of the chemotaxis plates were cleaned of condensation and debris using a dry cleaning wipe before placing them upright on the bed of an Epson V600 scanner without lids (S2 Fig). The plates were oriented such that quadrants QIV and QIII were located on the top-left and top-right, respectively, when viewed from above so the scanner path travels along the plate’s vertical axis. This results in the test quadrants (QI and QIV) appearing in the top-right and bottom-left of the resulting scanned images. A black paper backing was attached to the scanner background to improve worm contrast on the plates. TIF images were taken using the Epson Capture Software (Ver 3.9.3.1) at 1,200 dpi in greyscale with the Color Correction setting. This resolution was chosen as it optimally balanced accurate downstream processing with scanner capture speed. A custom MATLAB script was then used to process the images and calculate relevant behavioral metrics for each plate.

### Worm identification

The scanned TIF files were processed using a custom MATLAB algorithm (MATLAB 2023a). First, the plates were individually cropped out of the original scanner image by segmenting via pixel intensity and using the regionprops() function to map a circle to each object and isolate those with an equivalent diameter near 2,300 pixels (corresponding to the 5 cm plates) (S3 Fig). To account for minor differences in plate placement and angle on the scanner, each cropped plate image was translated and rotated such that the center dot of the plate became the center of the image and the “X’s” corresponding to the test compound sites were located in the top-right and bottom-left corners of the image. To identify worms within the re-oriented plate, the objects within the plate were filtered by size, pixel value, equivalent diameter, and Euler number. As worms that left the center region tended to clump differently depending on the proximity to other worms, we accounted for four different clumping scenarios that could affect the 2D area of the worm. While a single, linear individual worm is roughly 200 pixels (0.09 mm^2^), individual worms (Single) were defined as those with an area between 120-320 pixels (0.05-0.14 mm^2^) to accommodate for varied postures and size discrepancies. To account for varied clumping geometries, two (Double) or three (Triple) overlapping worms were defined as those between 320-550 (0.14-0.25 mm^2^) and 551-700 pixels (0.25-0.31 mm^2^), respectively. Worms that formed larger colonies (>3) tended to align into tight monolayer groups, so the number of worms within these larger clusters was estimated by dividing their total pixel area by the approximate size of one linear worm (200 pixels).

### Behavior quantification

To calculate the Chemotaxis Index (CI) of a plate, the centroids of each worm were mapped to a coordinate system and divided into either test or control regions. Worm centroids within 400 pixels (approximately 1 cm) from the center of the plate are considered non-mobile and excluded from these regions. The CI is then calculated according to Equation 1, where CI<0 indicates avoidance and CI>0 indicates attraction (Equation 1).

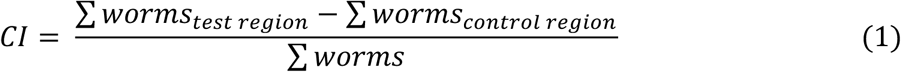

The Mobility of a plate is calculated as the proportion of worms that left the center region out of the total worms on the plate (Equation 2).

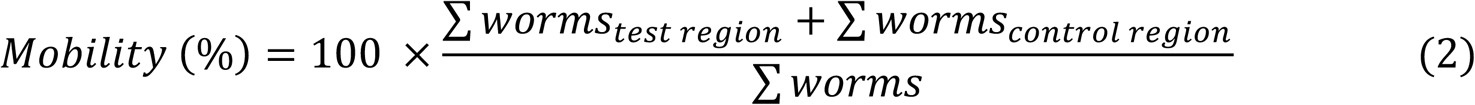

The Slope and Distribution of the worm Migration Path were calculated by mapping a linear model to the worm centroid coordinates and calculating its slope and adjusted R^2^ value. The Average Distance a worm travelled on each plate was found by taking the average of the distances of the worm centroids to the center of the plate, excluding those of non-mobile worms. For time-lapse datasets, this value was then used to calculate Average Movement Speed by dividing the Average Distance by the time elapsed between scanning intervals (5 min).

### Statistical analysis

All assays were performed with 3-5 technical replicates per condition for at least 3 biological replicates. All statistical analyses were performed in R (R Studio ver 4.3.1) using the stats (linear modeling, k-means clustering, ANOVA, t-tests, Pearson correlation tests) and DescTools (Dunnett’s test) packages. One- or Two-Way ANOVA was performed before post-hoc tests, assuming a significance threshold of α<0.05. Unless otherwise noted, asterisks indicate significance values determined by pairwise t-tests (two-tailed, unpaired, unequal variance) with Bonferroni correction (* p < 0.05, ** p < 0.01, *** p < 0.001).

## Results

### Development of an automated chemotaxis scoring platform

To increase the throughput and depth of *C. elegans* chemotaxis assays, we used the Lifespan Machine [39] strategy to capture worm positions within a plate using flatbed scanners and image-processing algorithms. We used an Epson V600 flatbed scanner and tested several physical parameters and image capture settings. To improve the contrast between the worms and agar in the plates, we added a black paper backing to the scanner lid (S2 Fig). Since the scanner can warp 3D features of objects placed furthest from the center of the scanner bed, leading to the walls of Petri dishes obstructing the viewable region of the plate, we restricted plate placement to 1-3 columns close to the center axis.

To identify *C. elegans* within images of scanned plates, we implemented a simple MATLAB algorithm adapted to grayscale images captured at 1,200 dpi, as these settings were optimal for balancing capture speed, worm detection accuracy, and processing time. The algorithm detects worms using pixel intensity, size, and Euler number and classifies them into one of four clumping patterns (see Methods). While our image-processing algorithm accommodates both round- and flat-bottomed Petri dishes, we recommend using the latter as the distorted edges of round-bottomed dishes reduced the total viewable region. We developed the platform using Margie et al.’s [18] quadrant-plate assay design with a 1 cm center region for initial worm placement (S3 Fig 3). The algorithm is designed to analyze plates with marks where worms and compounds are to be added, without the need for marking the quadrant boundaries, greatly reducing the amount of labor necessary to prepare the assay plates.

### Scanner platform validation

To validate the accuracy of TAXISCAN, we initially compared the results of the platform to those of standard manual scoring and found that, while the total number of worms it detected closely matched that of manual scoring, there were discrepancies (Pearson Correlation R = 0.88, 95% confidence interval 0.82 to 0.92, p<0.001). To probe the cause, we manually scored the same scanned plates in ImageJ and found that the results conflicted with those of standard scoring (S4 Fig). In particular, the largest disparity occurs when there are more than 150 worms on the plate. When comparing the number of worms detected by TAXISCAN against that of the manually scored images, there was very high agreement (Pearson Correlation R = 0.95, 95% confidence interval 0.93 to 0.97, p<0.001) (Figs 1A and S4 Fig) regardless of population size, suggesting that TAXISCAN can actually outperform standard (i.e., manual) scoring. In addition to detecting worms, TAXISCAN successfully enabled their localization relative to the plate geometry and calculate the Chemotaxis Index (CI) (Pearson Correlation R = 0.98, 95% confidence interval 0.97 to 0.99, p<0.001) (Fig 1B). Population size had no significant effect on CI accuracy (S5 Fig). We therefore recommend maintaining >100 worm population sizes on the assay plates, which is standard for chemotaxis assays to reduce the impact of natural behavior variability [2,3,18,28].

**Figure 1.**
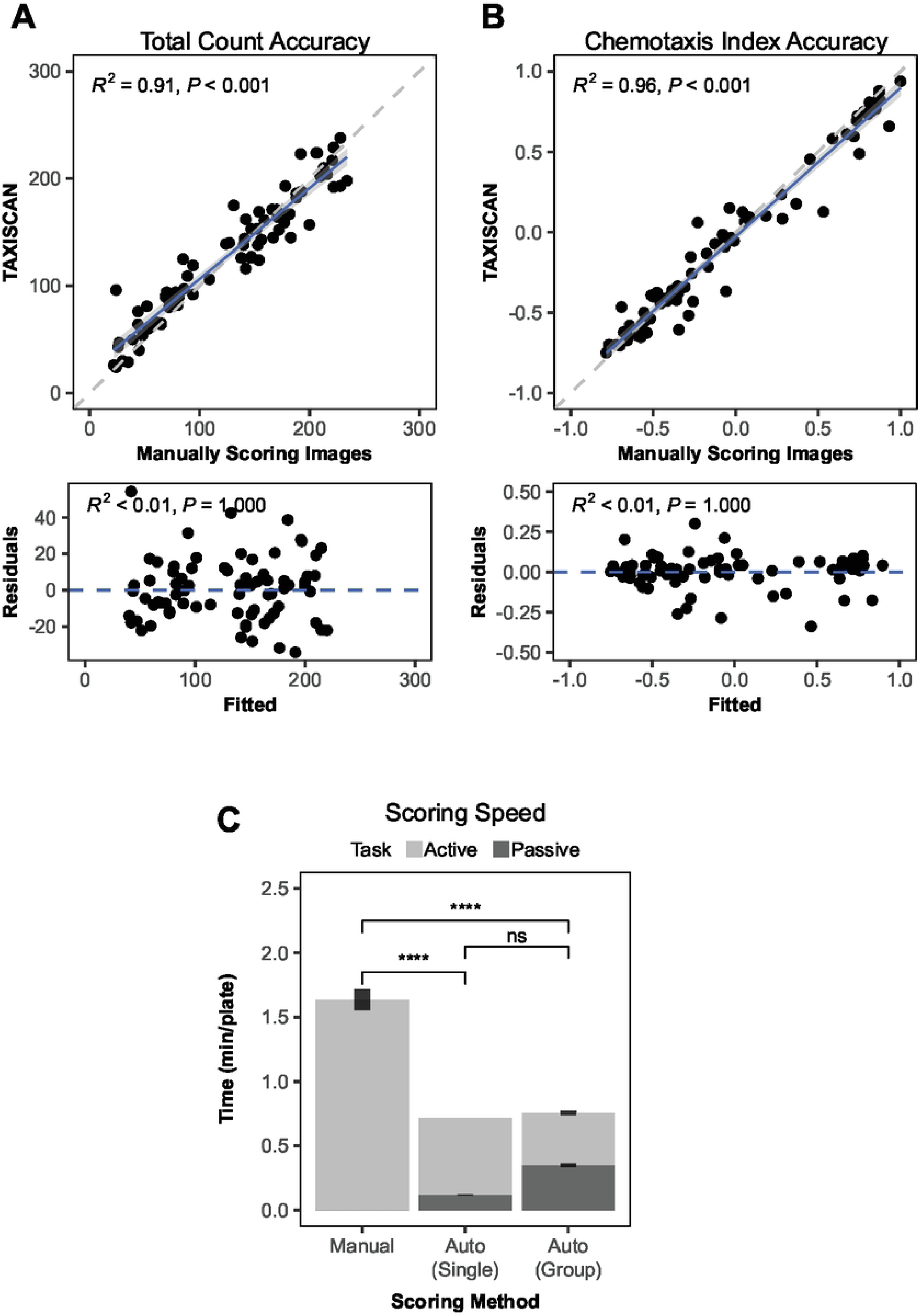
Performance of TAXISCAN. **(A-B)** Correlation and residual plots comparing the **(A)** total number of worms detected on a plate and the **(B)** resultant Chemotaxis Index for a chemotaxis plate scanned and scored by an observer (Manually Scoring Images) or the TAXISCAN system. The grey shaded region represents the 95% confidence interval for the fitted correlation model (dark blue line), plotted against an ideal correlation of slope=1 (grey dotted line). **(C)** The average time taken (per plate) for an observer to manually score a physical chemotaxis plate (Manual) compared to the time taken by TAXISCAN when processing plates one-by-one (Auto (Single)) or within a batch of multiple plates (Auto (Group)). The time taken for each task is further divided into steps requiring active participation (Active, light grey) such as scanning or manual scoring, versus those that can operate autonomously (Passive, dark grey) such as the image processing of TAXISCAN. Points represent the results of individual plates, and error bars are SEM.

To validate the utility of the platform, we compared the time it takes to fully score a plate (including both scanning and processing time) versus manual scoring. Scanning time depends on the number of plates along the scanner path; it takes roughly 40 s to scan a single 5 cm plate, 2 min for a column of up to 5 plates (24 s/plate), and 3 min for a 3×4 (15 s/plate) or 3×5 (12 s/plate) grid of plates. Compared to manual scoring, automation of a moderate chemotaxis assay (12-20 plates, 100-200 worms per plate) reduces total data acquisition time from 1.63 min per plate to 0.72 min when processed individually, or to 0.76 min when processing multiple plates at once (Dunnett’s test, p<0.001) (Fig 1C). While both automated methods take the same total time, processing multiple plates requires 12 s less to scan but 14 s longer to score per plate. Nonetheless, using the platform significantly reduces the active labor time from 1.63 min of manually scoring to 0.40 min of scanning, which would allow an experimenter to capture four times the replicates within the same time interval.

As *C. elegans* notoriously avoids light, we next tested if the light from the scanner impacted worm behavior and, if so, whether it was strong enough to override the natural chemotactic responses. To do so, we scored plates manually after the 60 min incubation period and then scanned them immediately. No significant movement or CI differences were observed between the manual counts scored before scanning and the algorithm-derived counts scored afterwards (1-way ANOVA, p== 0.98 and p= 0.6) (Fig 2A-B). We additionally tested the effects of the scanner light during the 60 min incubation period by repeatedly scanning the plates at 5 min intervals. We observed that, without chemical stimuli (Null, unpaired two-tailed t-test p=0.004), worms continuously exposed to the scanner’s light tended to prefer the bottom quadrants of the plates compared to unexposed worms that were scored manually (Figs 2A and 2C-D). This movement towards the bottom of the plates is consistent with avoiding the scanner light source, which is located in the back of the scanner or towards the top of the plates. However, this preference was not observed in worms exposed to attractants (IAA, unpaired two-tailed t-test p= 0.42) or repellents (CuSO_4_ and 1-octanol, unpaired two-tailed t-tests p= 0.96 and p= 0.16, respectively), and the overall CI was still unaffected for all compounds (1-way ANOVA p=0.98) (Fig 2B). Thus, maintaining the chemotaxis plates symmetrical along the scanner path, the effects of the scanner light on the behavioral responses to chemical stimuli are negligible, meaning TAXISCAN is a fast, accurate, and unobtrusive alternative to standard scoring techniques.

**Figure 2.**
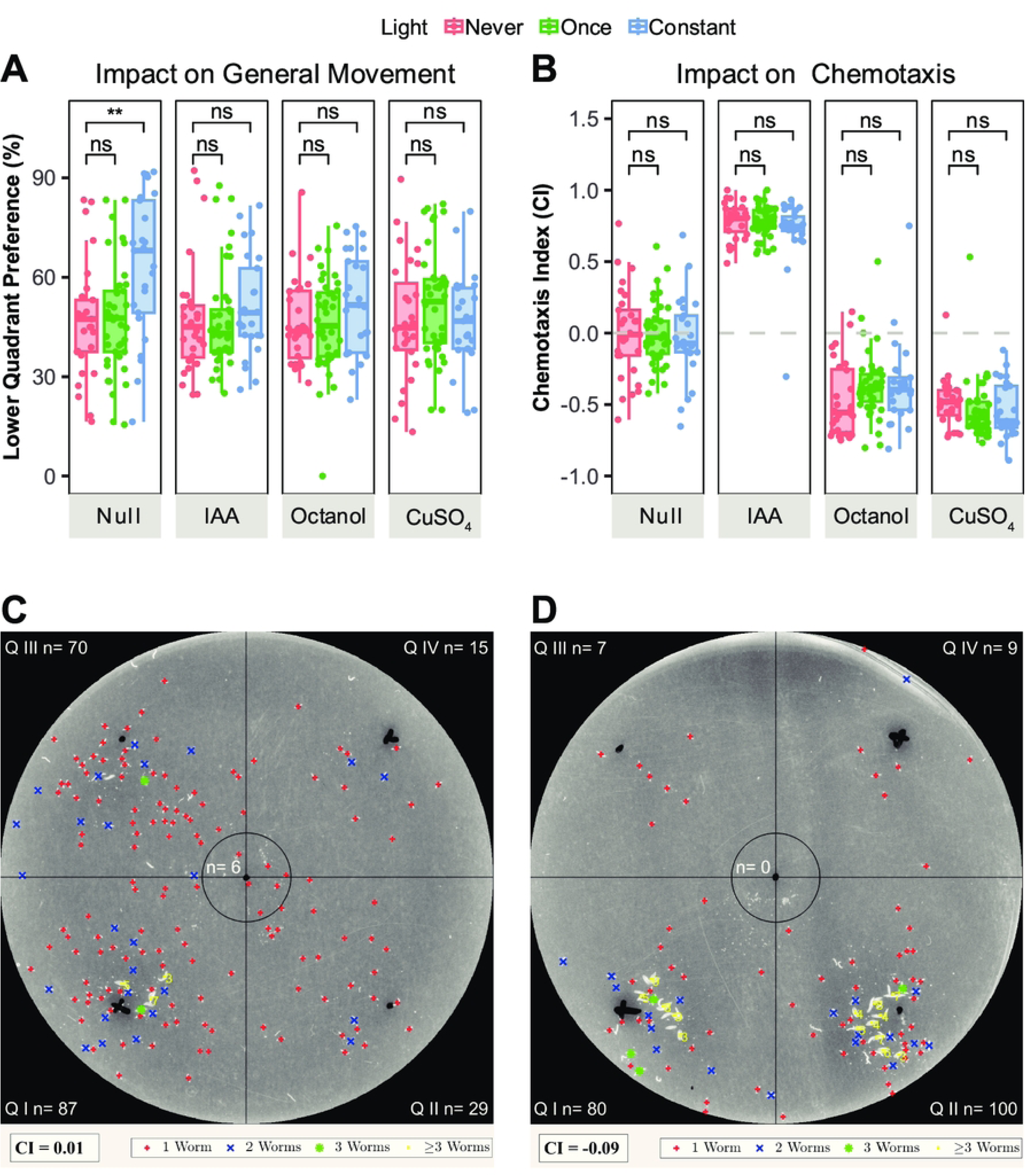
Impact of TAXISCAN on Behavior. The impact of exposure from the scanner light on **(A)** the percentage of mobile worms that moved towards the lower quadrants (QI & QII) of a chemotaxis plate and the resulting **(B)** Chemotaxis Index (CI) for the same plates, where CI>0 indicates attraction and CI<1 indicates avoidance. One cohort of plates were manually scored prior to light exposure (Never, red), then scanned and scored once by TAXISCAN (Once, green), while a second cohort of plates was continuously scanned every 5 min for 60 min (Constant, blue). Worms were exposed to 1% isoamyl alcohol (IAA) in 50% ethanol, 100 mM CuSO_4_, 10% 1-octanol in 50% ethanol, or chemotaxis buffer as a negative control (Null) with their respective buffers in the control quadrants of the plates. **(C-D)** Example plate readouts for negative controls after **(C)** one exposure to the scanner light or **(D)** repeated 5 min exposure during a 60 min assay. X’s denote where 2 μL of the test compound was added. Worms are labelled by whether one (red), two (blue), three (green), or multiple (≥3 yellow) worms were detected in a group.

### TAXISCAN reveals subtle chemotaxis behavior hidden in CI

As worm locations within a plate can be accurately mapped to a coordinate system, it is possible to characterize chemotaxis with additional behavior metrics (Fig 3A) such as travel distance and movement speed, which can help differentiate chemosensory from locomotion defects [24]. To test this principle, we used a *daf-19(m86)* chemosensory mutant (WBGene00000914) with no known locomotion defects but established chemosensory and roaming defects [40,41]. These chemosensory mutants exhibited reduced attraction to IAA (unpaired two-tailed t-test, p<0.001) and reduced avoidance to CuSO_4_ (unpaired two-tailed t-test, p<0.001), but still maintained typical avoidance to 1-octanol (unpaired two-tailed t-test, p=0.151) (Fig 3B). To determine if these differences were explained by defective chemosensation and not locomotion issues, we calculated the proportion of worms that left the center regions of the plates (Mobility) (Fig 3C) and how far they travelled (Migration Distance) (Fig 3D). Both wildtype and *daf-*19 mutants presented high Mobility (>80%) and similar Migration Distances (∼¼ of the plate in any direction, roughly 600 pixels or 12.71 mm) in response to all compounds but IAA. The reduced mobility in response to IAA is due to a behavioral defect, rather than a locomotory one, since the *daf-19* mutants travelled the same distance in response to IAA as for the other compounds (1-way ANOVA, p=0.25), which were all equivalent to that of wildtype (1-way ANOVA, p=0.25).

**Figure 3.**
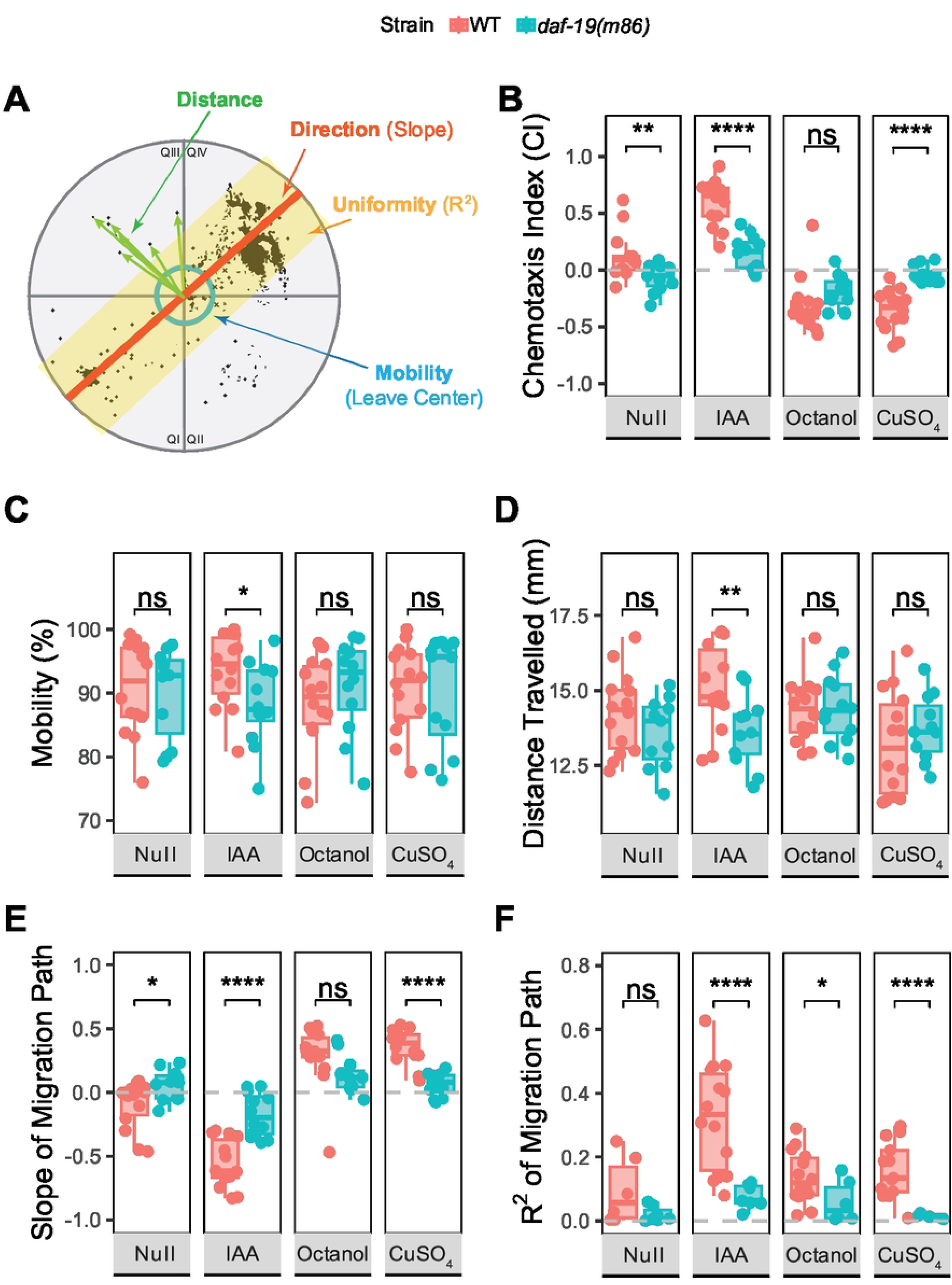
Capturing Multiple Behavioral Metrics. **(A)** Diagram showing how the Average Distance Travelled (green), Mobility (blue), Migration Slope (orange), and Migration Grouping (yellow) are calculated using an example plate readout for worms tested against 1% isoamyl alcohol (IAA). The plate is divided into quadrants (QI-QIV) where the test compound is located in QI and QIV, and the center region of the plate is outlined in black. The number of worms in this center region are excluded from quadrant counts except when referring to “total worms” on the plate. **(B-F)** Differences in chemotaxis behavior metrics between wildtype (WT, red) and *daf-19(m86)* mutant (blue) worms, including the **(B)** Chemotaxis Index (CI), **(B)** Percent Mobility, **(C)** Average Distance travelled (in pixels) by worms that left the center region, **(E)** Slope of the Migration Path, and **(F)** Variability in worm position about the Migration Path (R^2^). Worms were exposed to 1% IAA in 50% ethanol, 100 mM CuSO_4_, 10% 1-octanol in 50% ethanol, or chemotaxis buffer as a negative control (Null) with their respective buffers in the control quadrants of the plates.

To characterize subtler chemotaxis features, we examined the migration patterns within the plate by mapping the centroids of the detected worms to a linear model. The Slope of the model represents the Direction of the Migration Path (Direction) (Fig 3E), similar to a CI value, but accounts for the true positions of the worms instead of just the number of worms in each region. The R^2^ value reflects the general distribution of the worms about that path (Uniformity) (Fig 3F) and is impacted by worm grouping behaviors, which we found to vary by stimuli and population size (S6 Fig). In general, though, for wildtype worms a lack of chemical stimuli (buffer control, Null) results in a random distribution of worms from the center of the plate (Slope=0, R^2^<0.1), while attractants result in clumping at the compound site (Slope<0, R^2^>0.2), and repellent compounds result in diffuse distributions about a linear path directed away from the compound (Slope>0, 0.1<R^2^<0.2) (Fig 3E-F). We thus conclude that R^2^<0.1 indicates disorganized movement, R^2^>0.1 indicates voluntary and intentional movement, while R^2^>0.2 indicates highly focused and uniform movement. *daf-19* mutants do not exhibit this behavior, instead forming diffuse patterns across the plate regardless of the compound (R^2^<0.1, 1-way ANOVA p<0.001). This is also true for 1-octanol, which produced a similar CI value to wildtype (Fig 3B) but a significant difference in R^2^. Worms that form tighter clusters will exhibit a higher R^2^ than those that disperse evenly throughout a quadrant, despite having the same number of worms within the quadrant. Therefore, while the same proportion of wildtype worms and *daf-19* mutants avoided 1-octanol, the wildtype formed a more unified path to do so. This result suggests that although the CI metric does not identify a difference between wildtype and *daf-19* mutants, differences in chemotaxis behavior do exist but are only accessible through high-content metrics such as detailed distribution patterns (R^2^). Together, these findings show that TAXISCAN’s high-content analysis of chemotaxis based on quantitative worm position metrics reveal otherwise unidentifiable differences in chemosensory behavior.

The automated scoring pipeline also enables monitoring of CI and worm displacement throughout the course of a 60 min assay (Fig 4). At t=0 min, the majority of worms remain clustered in the center of the plate and prove difficult for the program to accurately score, but at t=5 min more than 50% of the worms have left the center region and are sufficiently dispersed to enable scoring (Fig 4A-B). In fact, across all compounds, worms travel a substantial fraction of the total distance within the first 5 minutes on the plate (Fig 4C), reaching a maximum speed of 2.11 mm/min, before rapidly decreasing to 0.37 mm/min, which steadily decelerates with time (Fig 4D). Because of this behavior, initial chemotaxis trends are visible at t=10 min (Fig 4A) and most prominent in the presence of an attractant (IAA), as the worms quickly leave the center of the plate and uniformly travel (R^2^>0.2) to the compound site within 20 min and remain there (Figs 4C, 4E-G, S7 Fig). Repellents, on the other hand, provoke slower and more nuanced responses. During 1-octanol stimulation, an initial, disorganized avoidant response (CI<0, R^2^<0.1) is visible at t=10 min but it takes up to t=30 min to observe a substantial uniform path (R^2^>0.2) (Figs 4A, 3E-F). CuSO_4_ likewise prompts avoidance within t=10 min, but it is evident through a diffuse path (R^2^>0.1) rather than haphazard positioning (Fig 4F), and the worms maintain this wide distribution as they proceed to the control quadrants at a slow but steady pace throughout the remainder of the 60 min observation (Figs 4A, 4E-F).

**Figure 4.**
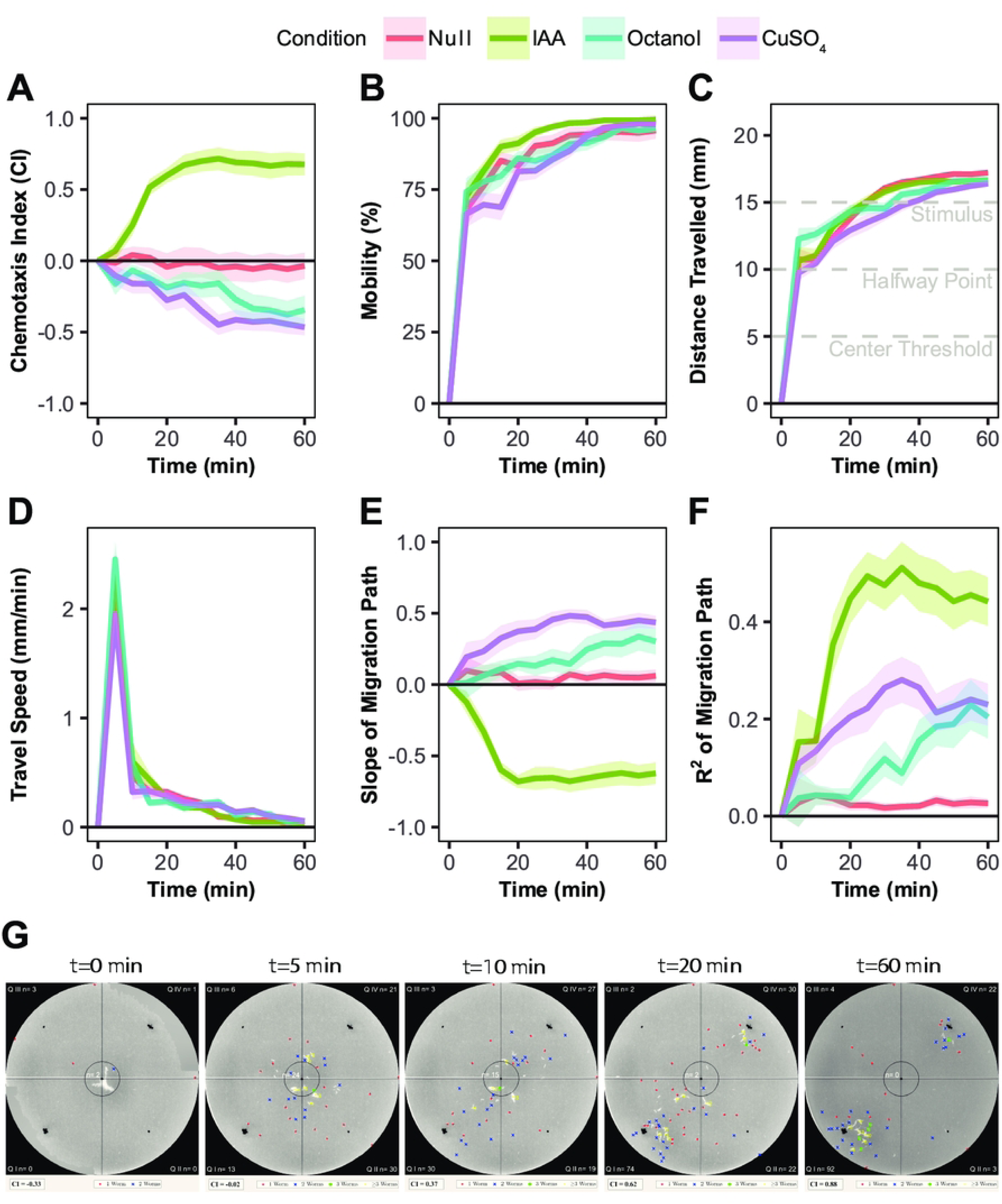
Dynamic Capture Reveals Distinct Behavior Patterns with Time. Time-lapse behavioral metrics observed in wildtype (N2) worms at 5 min intervals for a 60 min period. **(A)** Chemotaxis Index (CI), where CI>0 indicates attraction and CI<0 indicates avoidance. **(B)** The percentage of worms that left the 5 mm center region of the plate (Mobility). **(C)** Average Distance Travelled (in mm) and **(D)** Speed (in mm/min) of mobile worms that have left the 5 mm center region of the plate (Center Threshold). Distances are relative to the center of the plate, where the 2 μL of the stimulus is located at 15 mm (Stimulus). **(E)** Slope of the Migration Path, where slope>1 indicates avoidance and slope<1 indicates attraction. **(F)** Variability in worm position about the Migration Path (R^2^). Worms were exposed to 1% isoamyl alcohol (IAA) in 50% ethanol (green), 10% 1-octanol in 50% ethanol (blue), 100 mM CuSO_4_ (purple), or chemotaxis buffer as a negative control (Null, red). Shading around the lines is SEM. **(G)** CI and Time-lapse images of a plate with 1% isoamyl alcohol at the “X” marks in the top-right and bottom-left quadrants (QI & QIV) at t=0, 5, 10, 20, and 60 min. Worms were added to the dot at the center of the plate while the buffer control (50% ethanol) was added to the dots in quadrants QII and QIII. Worms are labelled by whether one (red), two (blue), three (green), or multiple (yellow) worms were detected in a group.

These patterns were also evident when mapping the collective movement of all worms over time (S7 Fig). Worms are known to exhibit a foraging behavior referred to as “random walk” when they are unable to detect local olfactory stimuli. This foraging behavior involves a roughly 10 min window of local search wherein the worm travels short distances due to rapid pirouetting, which is followed by an extended global search period wherein the worm travels longer distances between turns [42,42–44]. We were able to observe similar trends when mapping individual displacement over time; worms exposed to buffer controls remain close to the center of the plates during the first 10 min, consistent with local foraging behaviors. Then, at roughly 15 min, a bimodal response in worm positions is visible, reflecting the transition from local to global search. The shift from local to global search is accelerated by the presence of olfactory stimuli, especially volatile compounds (isoamyl alcohol & 1-octanol), but delayed in CuSO_4_, consistent with previous observations using Migration Path and Distribution (Fig 4E-F). While the presence of buffers and paralytics prevents these observations from truly reflecting a random walk, TAXISCAN can recapitulate this behavior typically observed via single-worm trackers quantifying pirouetting in addition to capturing population-level foraging activity. In summary, this automated chemotaxis platform enables characterization of different behavior patterns over time, even for chemicals that have similar CI values at the standard 60 min endpoint assay or for behaviors typically quantified via higher-resolution platforms. Moreover, this analysis revealed that attractants prompt rapid and uniform migrations while repellents prompt slower and more diffuse migrations that would otherwise be poorly captured by standard assays.

### TAXISCAN enables chemotactic analysis of mutants with altered locomotion

One substantial challenge for traditional chemotaxis assays is testing mutants with disrupted locomotion. Here, we sought to determine if the automated scoring platform could enable analysis of chemotactic responses in such challenging contexts, specifically assessing *slo-1* mutants as a test case. SLO-1 (WBGene00004830) is a potassium channel that affects neurotransmitter release at the neuromuscular junction (Fig 5A). s*lo-1* mutations are associated with perturbed neuronal activity, roaming behaviors, ethanol preference, and ethanol intoxication [45–49], but the effects of these mutations on chemotaxis behaviors outside of ethanol preference have not been well-studied. We tested three *slo-1* mutants including a loss-of-function with increased neuronal activity (LoF, *slo-1(js379)*), a gain-of-function with reduced neuronal activity (GoF, *slo-1(ky389)*), and a partial loss-of-function mutation (pLoF, *slo-1(gk602291)*) with resistance to ethanol-induced changes in locomotion and egg-laying [48,50], but uncharacterized effects on neuronal activity (Figs 5, S8 Fig, S9 Fig). Upon initial observations of the CI results, it appeared that all mutants had reduced IAA attraction, though the results of the other compounds were more nuanced; the high-neuronal activity LoF mutant presented severely reduced responses to all compounds while the low-neuronal activity GoF mutant merely exhibited slower CI responses (Fig 5B).

**Figure 5.**
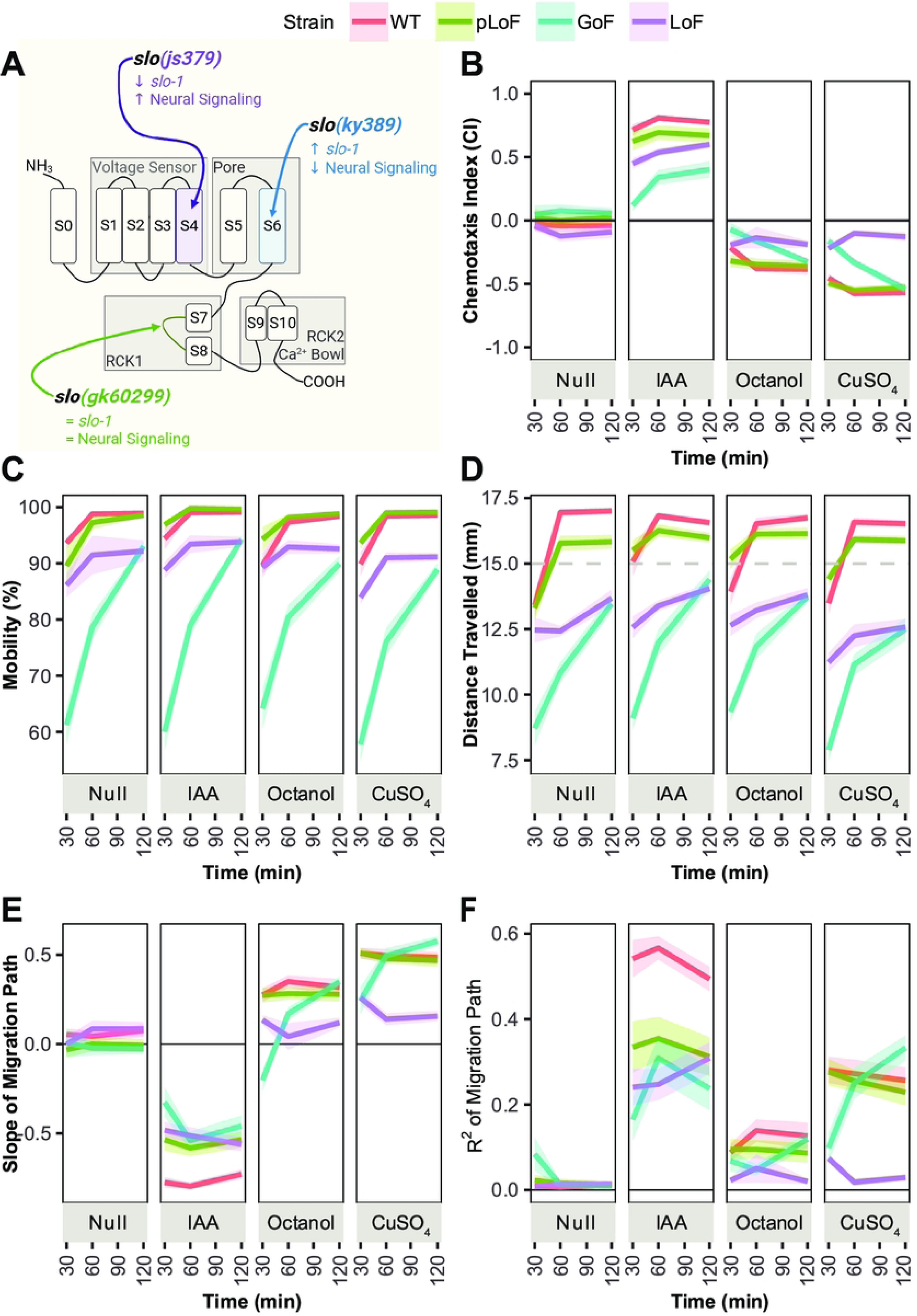
*slo-1* Mutations Impact Chemotaxis. **(A)** Graphic depicting the effect of different *slo-1* mutations on neuronal activity, based on graphics by Wang et al. 2001 [46] and Kokan et al. 2024 [50]. **(B-F)** Chemotaxis behavior metric readouts comparing wildtype (N2, red) worms to *slo-1(gk602291)* partial loss-of-function (pLoF, green), *slo-1(ky389)* gain-of-function (GoF, blue), and *slo-1(js379)* loss-of-function (LoF, purple) mutants. **(B)** Chemotaxis Index (CI), where CI>0 indicates attraction and CI<0 indicates avoidance. **(C)** The percentage of worms that left the 5 mm center region of the plate (Mobility). **(D)** Average Distance Travelled (in mm) of mobile worms that have left the 5 mm center region of the plate when 2 μL of the stimulus is located at 15 mm (grey dotted line). **(E)** Slope of the Migration Path, where slope>1 indicates avoidance and slope<1 indicates attraction. **(F)** Variability in worm position about the Migration Path (R2). Worms were exposed to 1% (IAA) in 50% ethanol, 10% 1-octanol in 50% ethanol, 100 mM CuSO_4_, or chemotaxis buffer as a negative control (Null). Shading around the lines is SEM. Some panels were created using BioRender assets.

The GoF mutant also exhibited significant locomotion deficits but only minor chemotactic deficits. While wildtype worms reached 80% mobility and traversed ¼ of the plate (∼600 pixels, ∼12.71 mm) within 30 min, it took 2 hr for the GoF mutants to do the same regardless of the compound (Fig 5C-D), indicating that they have a slow or lethargic phenotype. However, they still maintain relatively normal chemotactic ability, as after the 2 hr interval they matched wildtype CI, Slope, and R^2^ values for the repellents 1-octanol (Dunnett’s test, p=0.5) and CuSO_4_ (Dunnett’s test, p=0.89). Additionally, though their attraction to IAA is reduced (CI=0.26 vs 0.74, Dunnett’s test p<0.001), the Slope and R^2^ values of their Migration Path suggest that they are still weakly responding to this compound (Fig 5E-F). Thus, reducing neuronal activity via *slo-1* appears to affect locomotion without impacting chemosensation. These results are consistent with the known locomotion [48,51] and food-seeking behavior [52,53] defects associated with this GoF strain via its impact on muscle coordination and AWC activity, respectively.

In contrast to the GoF mutant, the LoF mutant exhibited only minor locomotion deficits but major chemosensory deficits. Compared to wildtype, it travelled roughly 3.18 mm (∼150 pixels) shorter distances and had lower Mobility (1-way ANOVA, p=<0.001) (Fig 5C-D), which was unexpected as this strain is associated with above-average movement speeds [49,54]. We additionally discovered an insensitivity to repellents; 1-octanol and CuSO_4_ both elicited weak CI values (−0.20 and −0.08, respectively) and nonresponsive R^2^ values (0.021 and 0.017, respectively), suggesting that the mutant was entirely unresponsive to both compounds. Although the LoF mutant also showed a reduced attraction to IAA (0.56 vs 0.74), its Slope and R^2^ values indicated uniform responses (Fig 5E-F). Therefore, the total LoF mutation has a significant impact on chemotactic ability, suggesting that increasing neuronal activity via *slo-1* can have an adverse effect on chemosensation. This conflicts with established observations that these mutations do not generally exhibit perturbed behaviors outside of ethanol intoxication [52,53]. However, chemosensory neurons are known to inhibit one another as part of the complex signal integration process [7,19,21,27,55,56], so it is possible that overactivity aggravates neuronal antagonism, leading to reduced chemotactic capacity.

From the three mutants, the chemosensory and locomotion ability of the pLoF mutant was the least affected. While a reduced attraction to IAA was observed (CI 0.56 vs 0.74), the Slope and R^2^ values indicated uniform responses (Fig 5E-F). The pLoF mutant exhibited no other chemotactic deficits but generally travelled 0.64 mm less than wildtype for all compounds (∼30 pixels) (Fig 5D). These results demonstrate that TAXISCAN can screen worms with challenging locomotion phenotypes and minor chemosensory deficits with ease and depth unfeasible in traditional assays.

## Discussion

In this work, we expand upon existing techniques for conducting standard chemotaxis assays through automated acquisition and scoring of chemotaxis plates in a rapid, accurate, and unbiased manner. Incorporation of a scanner dramatically reduces data acquisition time without significantly impacting worm behavior or requiring additional expensive equipment while improving accuracy of CI results. When paired with downstream image processing, this system allows for more replicates and increased sample size, which in turn can overcome the inherent variability associated with these behavioral assays. Additionally, as the exact positions of individual worms can be recorded in multiple plates simultaneously at multiple timepoints, TAXISCAN can be used to detect subtle behavioral differences through metrics such as Migration Distance, Direction, Uniformity, and Speed. These high-content metrics enable detailed analysis of the chemotactic behavior from severe *daf-19* chemosensory defects to the subtle altered locomotion and response delay phenotypes from *slo-1* mutations, of which the latter had been largely uncharacterized.

Our platform faces some limitations, as the accuracy of worm identification relies on high image quality as well as consistent worm sizes. The scanner bed and the chemotaxis plates have to be clean (i.e., no dust or scratches), and only a few plates can be scanned at a time as the scanner lens warps the plates furthest from the vertical center. Using short-walled, flat-bottomed plates with thin agar pads also produce clearer images compared to plates with >10 mL of agar or rounded edges. Additionally, the scoring platform is less accurate at distinguishing worms of abnormal size, those in large clumps, or those that have coiled into a ball or spiral. However, these issues can be addressed by the experimenter through manually amending the scoring algorithm’s parameters or correcting errors post-processing. Thus, despite its limitations, TAXISCAN poses a modular and accessible means to increase the throughput, accuracy, and depth of standard chemotaxis behavioral assays without the need for specialized equipment or training.

## Declaration of generative AI and AI-assisted technologies in the manuscript preparation process

During the preparation of this work, the authors used ResearchRabbit and Keenious Research Explorer to identify and cite literature relevant to the material presented. Otherwise, no generative AI tools were used in the design or execution of the material within this work. The authors have reviewed and edited the content as needed and take full responsibility for the content of the published article.

## Acknowledgements

*C. elegans* strains were provided by the *Caenorhabditis* Genetics Center (CGC). Funding for this project was provided in part by the NSF [2039226], the NCSU MBTP Training Program [T32GM133366], and the NCSU GAANN Fellowship. Some images in this work were created using BioRender assets. We would additionally like to thank the Dong Yan lab at Duke University (Durham, NC) for their assistance with developing the chemotaxis assay protocol.

## Author contributions

D.A.A. and V.R.Y. designed and implemented the image processing algorithm. V.R.Y. performed all experiments and analyses. V.R.Y. and A.S.M. interpreted results and wrote the manuscript.

## Supplemental

**Supplemental Figure 1. Chemotaxis Plate Layout.**

**(A)** Graphical representation depicting the design and layout of a 5 cm chemotaxis plate from a top-down view. **(B)** The same plate layout after processing using TAXISCAN, which captures plates from the bottom and crops the outer 0.5 cm border of the plate. Black X’s are marked 1 cm from the edge of the plate in quadrants QI (red) and QIV to correspond to 2 μL of the test compound (Orange). Small black dots are marked in the same spots of quadrants QII and QIII to denote the 2 μL of buffer control (blue), and the dot at the center of the plate denotes where worms are added. **(C-D)** Example worm distributions and Chemotaxis Indexes (CI) in response to **(B)** an attractant or **(C)** a repellent added in the test quadrants. The dark grey borders represent the edge of the 5 cm plate, and the light grey borders represent the 4 cm viewable window TAXISCAN crops out of the original image. The dotted lines represent the relative quadrant boundaries and the 1 cm diameter neutral region.

**Supplemental Figure 2. Scanner Setup.**

**(A)** Epson V600 scanner with a single column of chemotaxis plates. **(B)** The original white removable cushion on the back of the scanner lid, which was replaced with **(C)** a black paper backing to improve contrast in the resultant scanned images.

**Supplemental Figure 3. TAXISCAN Image Processing.**

Representative images demonstrating TAXISCAN’s processing of a column of chemotaxis plates. **(A)** A raw image captured using the Epson V600 scanner, taken from the bottom of the plates at 1200 dpi. **(B)** TAXISCAN’s plate recognition output, which defines the viewable region of the plates (red) and labels them (yellow text). **(C-E)** Downstream of a plate by **(C)** isolating and re-orienting, **(D)** segmenting the worms from the background, and **(E)** scoring by worm size and location. X’s denote where 2 μL of the compound was added, and worms are labelled by whether one (red), two (blue), three (green), or a large group (yellow) were detected. The plates are further divided into quadrants (black axes) and a 1 cm diameter neutral region (black circle), with *n* worms detected in each region (white). The Chemotaxis Index (CI) for the plate ranges between −1 and +1, representing total avoidance and attractance, respectively.

**Supplemental Figure 4. Accuracy in Worm Detection.**

Correlation and residual plots comparing the total number of worms detected on a plate by different scoring methods, clustered by if < 100 (red) or > 100 (blue) worms were detected on the plate. **(A, D)** Counting worms a plate under a microscope by hand (Standard Scoring) versus a scanned image of the same plate (Manually Scoring Images), with separate correlation models for each cluster. **(B-C, E-F)** Counting worms via TAXISCAN versus **(B, E)** Standard Scoring or **(C, F)** Manually Scoring Images. The correlation model (dark blue line) fit is given by the R^2^, p-value (P), and 95% confidence interval (grey shaded region), with slope=1 for reference (grey dashed line). Clustering by worm count was performed via k-means clustering.

**Supplemental Figure 5. Accuracy in Chemotaxis Index Determination.**

Correlation and residual plots comparing the Chemotaxis Index calculated for a plate by different scoring methods, clustered by if < 100 (red) or > 100 (blue) worms were detected on the plate. **(A, D)** Counting worms on a plate under a microscope by hand (Standard Scoring) versus a scanned image of the same plate (Manually Scoring Images), with separate correlation models for each cluster. **(B-C, E-F)** Counting worms via TAXISCAN versus **(B, E)** Standard Scoring or **(C, F)** Manually Scoring Images. The correlation model (dark blue line) fit is given by the R^2^, p-value (P), and 95% confidence interval (grey shaded region), with slope=1 for reference (grey dashed line). Clustering by worm count was performed via k-means clustering.

**Supplemental Figure 6. Impacts of Chemical Stimuli and Population Size on Worm Grouping Patterns.**

The percentage of wildtype (N2) worms that formed groups of 3 or more out of the total number of worms detected on a chemotaxis plate, sorted by **(A)** small and **(B)** large plate populations. Worms were exposed to 1% IAA (green) 10% 1-octanol in 50% ethanol in 50% ethanol (blue), 100 mM CuSO_4_ (purple), or chemotaxis buffer as a negative control (Null, red) with their respective buffers in the control quadrants of the plates. Error bars are SEM. Plate population sizes are separated into <100 (red) or >100 (blue) worms per plate, determined via k-means clustering.

**Supplemental Figure 7. Wildtype Migration Over Time.**

The kernel density estimate (Worm Density, left y-axis) for individual worm travel distances (Distances, x-axis) over time (right y-axis). Worms were exposed to 1% isoamyl alcohol (IAA, green) in 50% ethanol, 10% 1-octanol (blue) in 50% ethanol, 100 mM CuSO_4_ (purple), or chemotaxis buffer as a negative control (Null, red). Vertical lines indicate the approximate means of the bimodal peaks detected via standard EM algorithm. The center of the plate is located at x = 0 mm, and the compound is located at x = 15 mm.

**Supplemental Figure 8. s*l*o*-1* Mutant Migration Over Time.**

The kernel density estimate (Worm Density, y-axis) of the percentage of individual *slo-1* mutant worm travel distances from the center of the plate (Distances, x-axis) over time in response to chemotaxis buffer alone. Chemotaxis behavior metric readouts comparing wildtype (N2, red) worms to *slo-1(ky389)* gain-of-function (GoF, green), *slo-1(js379)* loss-of-function (LoF, blue), and *slo-1(gk602291)* partial loss-of-function (pLoF, purple) mutants. Vertical lines indicate the means of the bimodal peaks detected via standard EM algorithm. The center of the plate is located at x = 0 mm, and the compound is located at x = 15 mm.

**Supplemental Figure 9. s*l*o*-1* Mutant Chemotaxis.**

Representative chemotaxis images for Wildtype (N2, red) worms and *slo-1* partial loss-of-function (*slo-1(gk602291)*, green), gain-of-function (*slo-1(ky389gf)*, blue), and loss-of-function (*slo-1(js379lf)*, purple) mutants. Worms were exposed to 1% isoamyl alcohol (IAA) in 50% ethanol, 10% 1-octanol (Octanol) in 50% ethanol, 100 mM CuSO_4_, or chemotaxis buffer as a negative control (Null). X’s denote where 2 μL of the compound was added, and worms are labelled by whether one (red), two (blue), three (green), or multiple (≥3 worms, yellow) worms were detected in a group.

